# Enterococcal Polysaccharide Antigen (EPA) rhamnan backbone contributes to cell wall architecture and is essential for antimicrobial resistance, innate immune evasion and phage infection

**DOI:** 10.64898/2026.08.24.746643

**Authors:** Lea Kupcova, Nishan Nathoo, Bartosz J. Michno, Krishna S. Chellappa, Thomas Lawson, Mark McNeil, Jessica L. Davis, Pavithra Manivannan, Joshua S. Norwood, Robert E. Smith, Emmanuel Maes, Laia Pasquina-Lemonche, Tomasz K. Prajsnar, Michelle L. Rowe, Helge C. Dorfmueller, Graham Stafford, Mike P. Williamson, Stéphane Mesnage

## Abstract

Enterococci are opportunistic pathogens classified by the World Health Organization as high-priority microorganisms. They cause a broad spectrum of infections, and their intrinsic and acquired resistance to antimicrobials makes these infections particularly difficult to treat and eradicate. In *Enterococcus faecalis*, the most frequently isolated enterococcal pathogen in humans, antimicrobial resistance and innate immune evasion are largely driven by the Enterococcal Polysaccharide Antigen (EPA). This surface polymer underpins key virulence traits, including resistance to host defence mechanisms, reduced susceptibility to multiple classes of antimicrobials, and susceptibility to bacteriophage infection. EPA consists of a rhamnan backbone decorated with strain-specific substituents that are essential for its biological activity. Here, we show that epaB encodes the enzyme responsible for the first committed step in assembling the EPA rhamnan chain. Using NMR spectroscopy, we demonstrate that *E. faecalis* lacking epaB produces an EPA polymer composed solely of decorations directly anchored to the peptidoglycan, with no detectable rhamnan backbone. The absence of this rhamnan moiety profoundly alters cell wall architecture, as revealed by atomic force microscopy of the mutant cell walls. The epaB mutation also abolishes innate immune evasion and virulence in the zebrafish infection model, while conferring resistance to bacteriophages. Collectively, these findings demonstrate that both the rhamnan backbone and its decorations are required for EPA’s full biological activity, establishing the structural and functional interdependence of these two components.

**IMPORTANCE:** The Enterococcal Polysaccharide Antigen (EPA) is essential for normal growth and division, virulence, antimicrobial resistance, and phage infection in enterococci. This surface polymer comprises a structurally conserved rhamnan backbone substituted with strain-specific decorations. These variable decorations have been directly linked to the biological functions of EPA, whereas the rhamnan backbone has been proposed to serve primarily as a structural scaffold. Here, we use NMR spectroscopy to show that mutation of *epaB*, the gene responsible for the first biosynthetic step in rhamnan backbone formation, results in an EPA polymer composed exclusively of decorations anchored to the peptidoglycan. We reveal that the lack of rhamnan backbone is associated with a change in the cell wall architecture and abolishes virulence and infection by bacteriophages. Together, these findings demonstrate that assembly of the conserved rhamnan backbone is indispensable for EPA function and highlight this biosynthetic step as a promising target to combat enterococcal infections.

## INTRODUCTION

Enterococci are commensal bacteria that colonize the gastrointestinal tract of all land animals (1) Due to its intrinsic resistance to antibiotics, *Enterococcus faecalis* has emerged as a nosocomial pathogen frequently associated with hospital- and community-acquired infections (2). *E. faecalis* can cause life-threatening infections such as bacteraemia and endocarditis and the virulence of this organism has been extensively studied. A major and ubiquitous *E. faecalis* virulence factor is the Enterococcal Polysaccharide Antigen (EPA), a rhamnose-containing polysaccharide with a functional equivalent in other pathogenic ovo*cocci* such as *Streptococcus pyogenes* or *Streptococcus mutans* as well as in *lactococci* (3). The *epa* locus was originally identified in *E. faecalis* OG1RF through the screening of genomic expression libraries with enterococcal endocarditis patient sera, alongside specific antisera raised in rabbits against surface proteins from an endocarditis isolate (4). Immunoscreening allowed the identification of antigenic clones that reacted with patient sera, leading to the discovery of genes encoding EPA. This complex polysaccharide is encoded by two adjacent loci (5) (Fig. 1A). The first locus (*epaA-epaR*) is conserved across all *E. faecalis* strains and is responsible for the production of a rhamnan polysaccharide substituted by glucose (Glc) and *N*-acetylglucosamine (GlcNAc) residues (6). The second *epa* locus contains a variable number of genes that encode strain-specific “decorations” polymerised by glycerol phosphate (strain OG1RF) or ribitol phosphate groups (strain V583). EPA decorations play an essential role in biofilm formation, resistance to antimicrobials, phagocyte evasion (7–9) and recognition by bacteriophages (10, 11). The role of the EPA rhamnan backbone remains elusive.

**Fig. 1.**
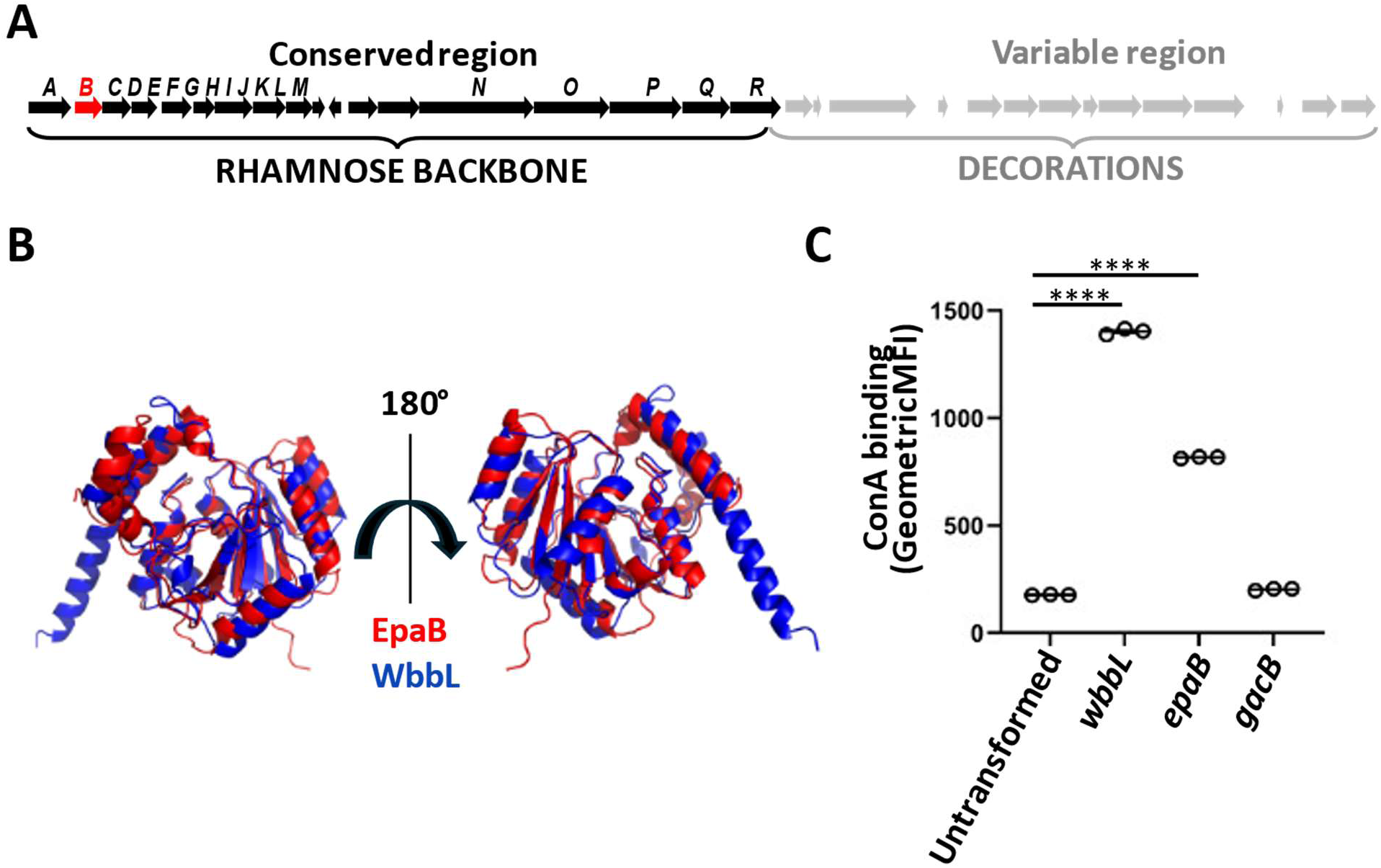
Restoration of *E coli* K-12 O-Antigen by the EpaB α-Rha-(1→3)-α-GlcNAc transferase. Detection of O-antigen assembly was achieved using Concanavalin A conjugated to Alexa Fluor 633. *E coli* K-12 Δ*wbbL* cells were transformed with plasmids encoding *E. coli* WbbL, *E. faecalis* EpaB and *Streptococcus pyogenes* GacB. Cells were stained with Concanavalin A conjugated to Alexa Fluor 633 and surface-associated fluorescence was analysed using flow cytometry. Geometric mean fluorescence intensities were compared using a one-way ANOVA and found significantly higher to untransformed cells when *E. coli* Δ*wbbL* cells were complemented with *wbbL* or *epaB* (p<0.0001; ****)

One of the EPA biosynthetic genes is *epaB*, a putative rhamnosyltransferase predicted to anchor the first rhamnose residue of EPA to the linkage unit attached to peptidoglycan. Phenotypic characterization of the *epaB* transposon mutant first reported (12) has been described in several studies, which identified defects in virulence or increased resistance to antimicrobials (13–15). Importantly, as these studies did not include complemented strains, polar effects could not be excluded. More recent work has described the in-frame deletion of *epaB* and its complemented derivative (16, 17), confirming that this mutation confers resistance to bacteriolytic agents (16) and impairs both biofilm formation and colitogenic activity in a mouse model (17). Both in-frame deletion, as well as the original mutation resulting from transposon mutagenesis were described as mutants unable to produce the EPA polymer (16–18).

Here we investigate the structure of EPA_B polysaccharide produced by the in-frame *epaB* deletion mutant (17). We demonstrate that EpaB is a rhamnosyltransferase that links the first residue of the rhamnan chain to the α-GlcNAc residue linked to peptidoglycan and show that the *epaB* mutation leads to the production of an EPA polymer devoid of rhamnose and directly anchored to peptidoglycan. The lack of a rhamnan chain in EPA leads to a defect in the cell wall architecture, resistance to bacteriophages and abolishes phagocyte evasion in the zebrafish model of infection. Collectively, these results indicate that although EPA decorations are essential for the biological activity of this surface polymer, they are not sufficient and require the presence of the rhamnan chain.

## RESULTS

### EpaB displays α-Rha-(1→3)-α-GlcNAc transferase activity and restores O-antigen synthesis in *E. coli*

The gene encoding *epaB* is located at the 5’ end of the *epa* operon (Fig. 1A) and encodes a putative glycosyltransferase. Foldseek predictions indicate that EpaB is structurally related to WbbL α-Rha-(1→3)-α-GlcNAc-pyrophosphoryl-undecaprenol transferases characterised in *Mycobacterium tuberculosis* and *E. coli* (19, 20). Despite a relatively low sequence identity (less than 26% for both homologs), structural predictions revealed that *E. faecalis* EpaB is most similar to *E. coli* WbbL, with an RMSD value of 2.36Å over 228 residues; Fig. 1B).

To investigate whether enterococcal EpaB can functionally substitute for WbbL, we expressed *epaB* in a Keio collection Δ*wbbL* K-12 *E. coli* mutant and assessed O-antigen production by Concanavalin A (ConA) staining followed by flow cytometric analysis (Fig. 1C) (21). Expression of *epaB* restored ConA staining to levels substantially above the Δ*wbbL* background control, indicating successful reconstitution of O-antigen biosynthesis. The positive-control strain expressing *E. coli wbbL* restored O-antigen production, whereas the Δ*wbbL* strain carrying empty vector remained ConA-negative. Notably, GacB, an α-D-GlcNAc-β-(1→4)-rhamnosyltransferase, was unable to complement the Δ*wbbL* mutant despite catalysing a related rhamnosyltransferase reaction (22). This finding suggests that restoration of O-antigen biosynthesis requires the specific α-(1→3) rhamnosyltransferase activity provided by EpaB. Together, these results demonstrate that EpaB functions as a WbbL homologue *in vivo* and is sufficient to restore O-antigen biosynthesis in a Δ*wbbL E. coli* strain.

### Solution-state NMR reveals that EPA_B, produced by the Δ*epaB* mutant, lacks the rhamnan backbone and only contains terminal decorations

Cell walls from the Δ*epaB* mutant were purified and digested with mutanolysin to solubilise intact peptidoglycan-anchored polymers. Following size-exclusion and ion-exchange purification, the material was analysed using solution state nuclear magnetic resonance (NMR). Phosphorus NMR (^31^P NMR) revealed the presence of all signals corresponding to EPA decorations (Fig. S2) whilst ^1^H signals lacked methyl protons (between 1-1.5 ppm) characteristic of rhamnose residues (Fig 2A). ^1^H-^13^C HSQC NMR confirmed the absence of all anomeric signals previously assigned to the rhamnan backbone (Fig. 2B). A total of 22 spin systems, named A-Q were identified, 5 of which are alternative variants of other spin systems with different substitution patterns (named B’, I’, M’, P’, and O’). Using the strategy previously described (11), we assigned all signals detected in the HSQC spectra (Table S1, Table S2, Fig. S3 and characterised residue connectivity (Table S3). The structural analysis showed that EPA_B decorations are indistinguishable from those present in the parental strain. Based on the integration of signal intensities, the polymerisation of the decorations is similar to the structure of EPA_WT elucidated by our group previously and contains 8-10 repeats.

**Fig 2.**
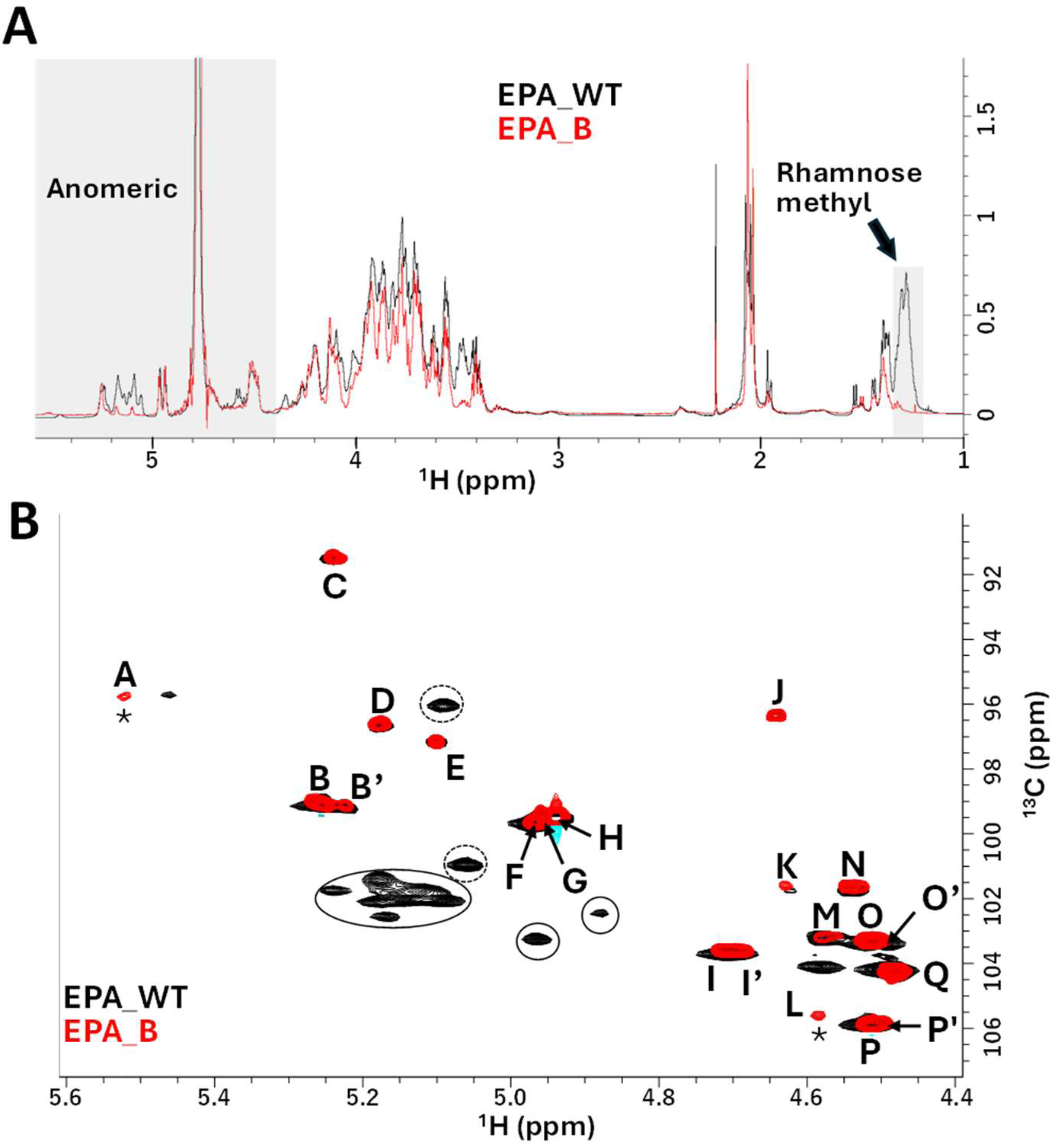
Solution state NMR reveals that EPA_B lacks rhamnose residues. (**A**) ^1^H NMR signals corresponding to methyl protons characteristic of rhamnose residues in *E. faecalis* OG1RF wild type (EPA_WT, black) are absent in *E. faecalis* OG1RF Δ*epaB* (EPA_B, red). Several signals in the anomeric region (grey) are also missing in the EPA_B polysaccharide. (**B**) ^1^H-^13^C HSQC spectra confirm that WT rhamnose signals (circles) as well as Glc and GlcNAc modifying the rhamnan chain (dotted circles) are absent in EPA_B. The signals corresponding to the residues previously identified in EPA_WT decorations are still present in EPA_B (11), except for two signals (asterisks) with no equivalent in EPA_WT.

Apart from the rhamnose signals, only two noticeable differences were found in the anomeric region of EPA_B (indicated by asterisks in Fig. 2B). The signal at 4.58 ppm was found to correspond to a low abundance β-glucose residue (labelled L in Figure 2B) that cannot be defined as part of the structure due to a lack of NOE signals. The signal corresponding to the GlcNAc residue A linked to the peptidoglycan MurNAc sugar in EPA_WT (5.46 ppm) was shifted to 5.52 ppm in EPA_B. As expected, ^1^H-^31^P HSQC and ^1^H-^31^P HSQC-TOCSY confirmed that residue A is a GlcNAc corresponding to the linkage unit, bound to position 6 of MurNAc via a phosphate bond (Fig. 3A). The analysis of NOESY spectra revealed a NOE peak from residue M to the same proton shift as a peak in residue A (Fig. 3B). The link between residues A and M was determined to be a 1-4 glycosidic bond using COSY cross peaks (Fig. 3C). Collectively, our NMR analysis showed that the structure of EPA_B corresponds to WT decorations anchored to a GlcNAc residue attached to the peptidoglycan MurNAc residue via a phosphodiester bond (Fig. 3D).

**Fig 3.**
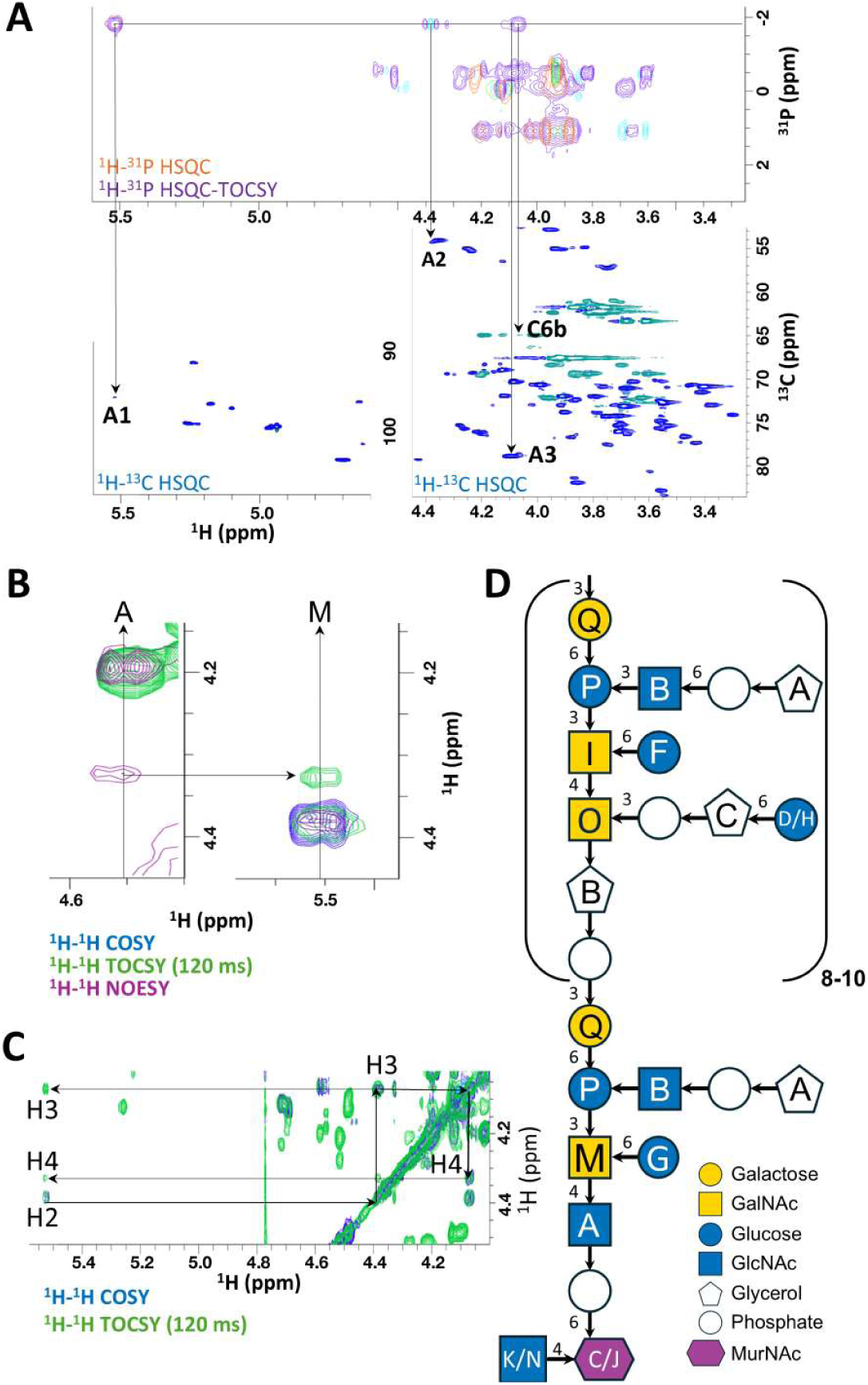
EPA_B consists of decorations linked directly to the peptidoglycan. (**A**) Residue A (α-GlcNAc) is linked to position 6 of residue C (α/β-MurNAc, from the peptidoglycan) through a phosphate bond. Peaks at −1.83 ppm in the ^1^H-^31^P HSQC (orange) and ^1^H-^31^P HSQC-TOCSY (purple, top panel) map to peaks corresponding to positions 1 (anomeric), 2, and 3 of A (blue peaks), and position 6 of C (green anti-phase peaks) in the ^1^H-^13^C HSQC spectra (bottom two panels). **(B**) A ^1^H-^1^H NOESY peak (pink) visible at 4.33 ppm from residue A (α-GlcNAc) corresponds to the same chemical shift as a ^1^H-^1^H TOCSY shift (green) of residue M (β-GalNAc). **(C**) Based on ^1^H-^1^H COSY cross peaks (blue) matching the ^1^H-^1^H TOCSY peaks (green) at the anomeric shift of A, the peak at 4.33 ppm was determined to arise from position H4 of A. indicating that residues A and M are linked through a 1,4 glycosidic bond. **(D**) Complete structure of EPA from *E. faecalis* OG1RF Δ*epaB* (EPA_B). Labelling of sugars matches labelling in Figure 2B.

### Solid-state NMR indicates that the lack of a rhamnan chain alters the surface exposure of EPA

The impact of the *epaB* mutation on EPA surface display was further investigated by solid-state NMR. We used High-resolution magic angle spinning (HR-MAS) NMR, to compare the solvent accessibility of EPA residues in OG1RF and its Δ*epaB* isogenic mutant (Fig. 4). Comparative analysis of HR-MAS spectra revealed reduced signal intensities for several resonances corresponding to decoration residues in the Δ*epaB* mutant (I, M and P). These findings suggest that the rhamnan chain plays a role in promoting surface exposure of EPA decorations.

**Fig 4.**
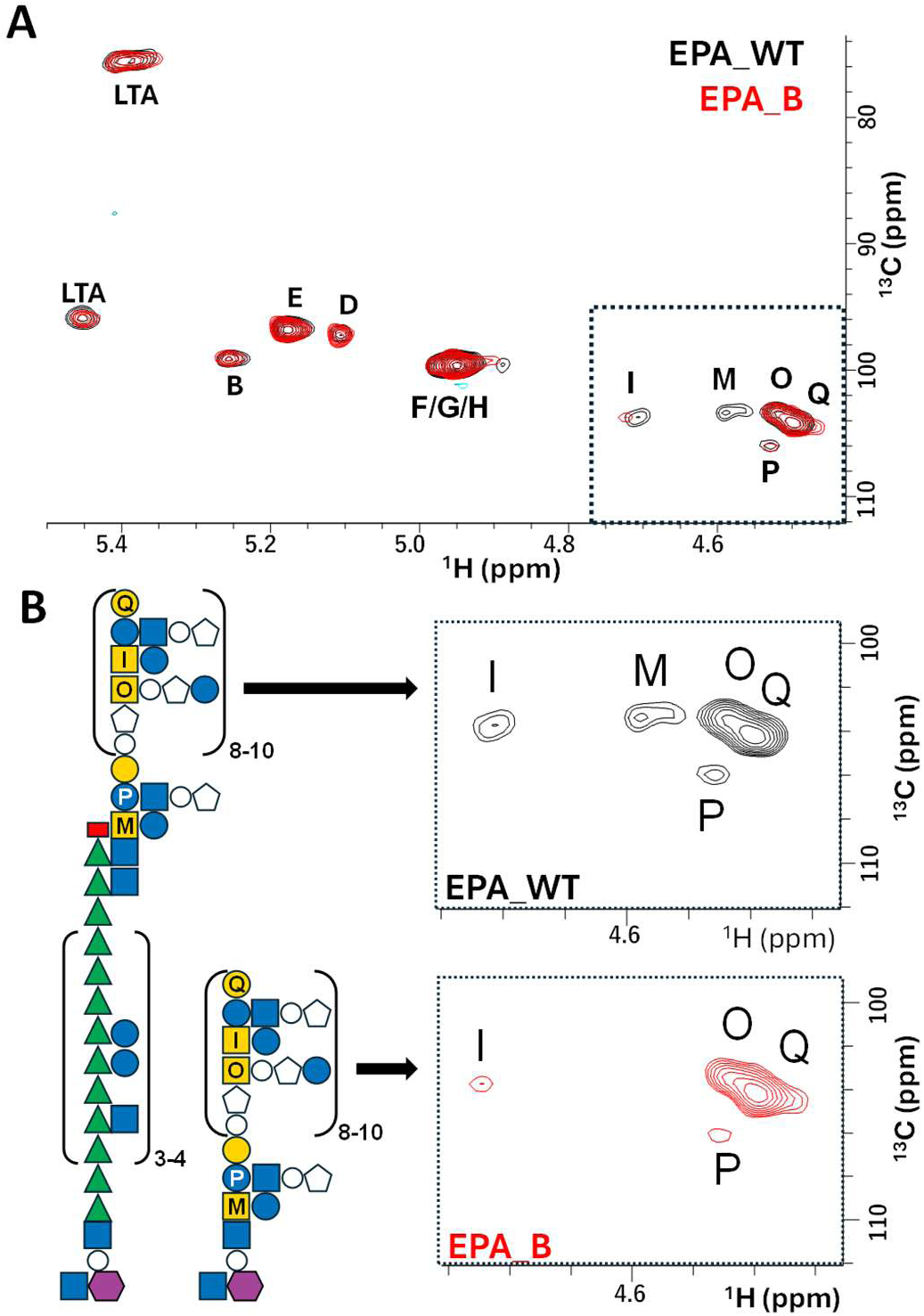
Solid state NMR ^1^H-^13^C HSQC HR-MAS analysis of *E. faecalis* OG1RF and Δ*epaB*. (**A**) Intensities from spectra recorded on *E. faecalis* OG1RF wild-type (EPA_WT, black) and *E. faecalis* OG1RF Δ*epaB* (EPA_B, red) were adjusted by matching to the signal intensities corresponding to lipoteichoic acids (LTA). **(B**) Comparison of signals corresponding to EPA revealed that residues I, M and P decrease in intensity from EPA_WT (black) compared to EPA_B (red), indicating that in the absence of the rhamnan, decorations are less solvent exposed. The methyl cap group is represented by a red rectangle.

### *epaB* mutants are resistant to infection by bacteriophages

We previously described several lytic bacteriophages that require EPA decorations to cause infection (11). Three phages requiring distinct EPA decoration motifs for optimal infections were chosen (Fig. 5A) to test how the lack of rhamnan backbone impacts phage susceptibility using efficiency of plating as a readout. Despite the presence of structural motifs required for phage infection in the EPA_B polymer, all three phages displayed a lower infectivity against the Δ*epaB* mutant as compared to the WT. Efficiency of infection was restored in all cases when phages were tested against the complemented Δ*epaB* strain (Δ*epaB + epaB*). These results agree with HR-MAS NMR suggesting that the surface exposure of decorations is altered when the rhamnan chain is not part of EPA.

**Fig 5.**
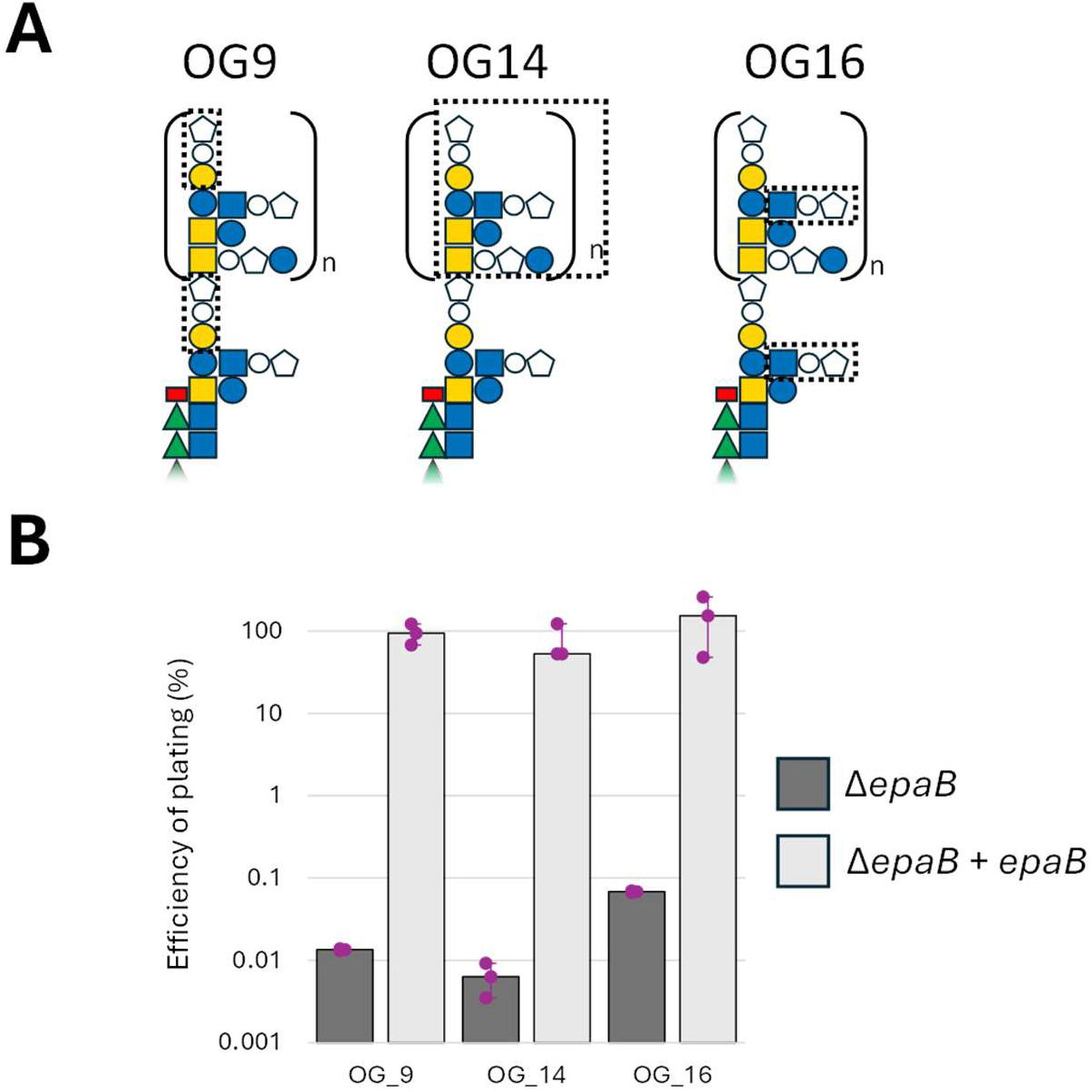
*E. faecalis ΔepaB* is resistant to bacteriophage infection. (**A**) Dashed boxes indicate the EPA structural motifs critical for infection by phages OG9, OG14 and OG16. (**B**) Efficiency of plating of phages against *E. faecalis* (*ΔepaB*, dark grey) and complemented derivative (*ΔepaB+epaB*, light grey). Standard deviation is indicated as purple error bars.

### The lack of rhamnan in EPA alters cell surface architecture

HR-MAS results and phage susceptibility assays suggested that the lack of the rhamnan chain alters the enterococcal cell wall architecture. Cell wall extractions from cultures gave similar mass yields for the WT, Δ*epaB* mutant and complemented counterpart (Fig. 6A). In all strains, the proportion of peptidoglycan and polymers in all strains remained unchanged (no statistical differences were found), accounting for approximately 30% of the cell wall dry weight (Fig. 6A). Atomic force microscopy (AFM) imaging of purified sacculi (containing both peptidoglycan and EPA) in air indicated that Δ*epaB* cell walls in a dry environment were significantly thinner than the WT (Fig. 6B). High resolution AFM imaging of intact, whole cells in a liquid environment (23) revealed obvious changes in overall bacterial cell morphology (Fig. 6C-D (i)) as well as nanoscale differences in the cell wall architecture in the Δ*epaB* mutant (Fig. 6C-D (ii-iii)). We used the custom-made open-source AFMSlicer FIJI routine (https://zenodo.org/records/13133978), which slices the AFM images in 2D binary slices along the Z axis to measure the abundance and geometry of cell wall pores at certain regions of interest. We established that OG1RF WT and its Δ*epaB* derivative have similar porosities (as defined by the ratio of material to empty space) (Fig. 6E (i)) alongside similar average pore diameter (Fig. 6E (ii)). However, significant differences were found between the cross-sectional shape of the pores (Fig. 6E (iii)) and their depth into the cell wall (Fig. 6E (iv)). The WT cell wall exhibits significantly more cylindrical pores with a slope of 1.021 when plotting the representative area against slice depth. In Δ*epaB* cell walls, that slope corresponded to more conical pores with a value of 1.953. The pore depth of the region of interest analysed indicates that those cylindrical pores in the WT penetrates 6.54 nm into the cell wall, whereas the conical pores in the cell wall of the Δ*epaB* mutant only penetrate 3.57 nm into the wall, resulting in a significant difference in pore depth. Collectively, these results indicate that the composition of EPA plays a key role in the architecture of the enterococcal cell wall.

**Fig 6.**
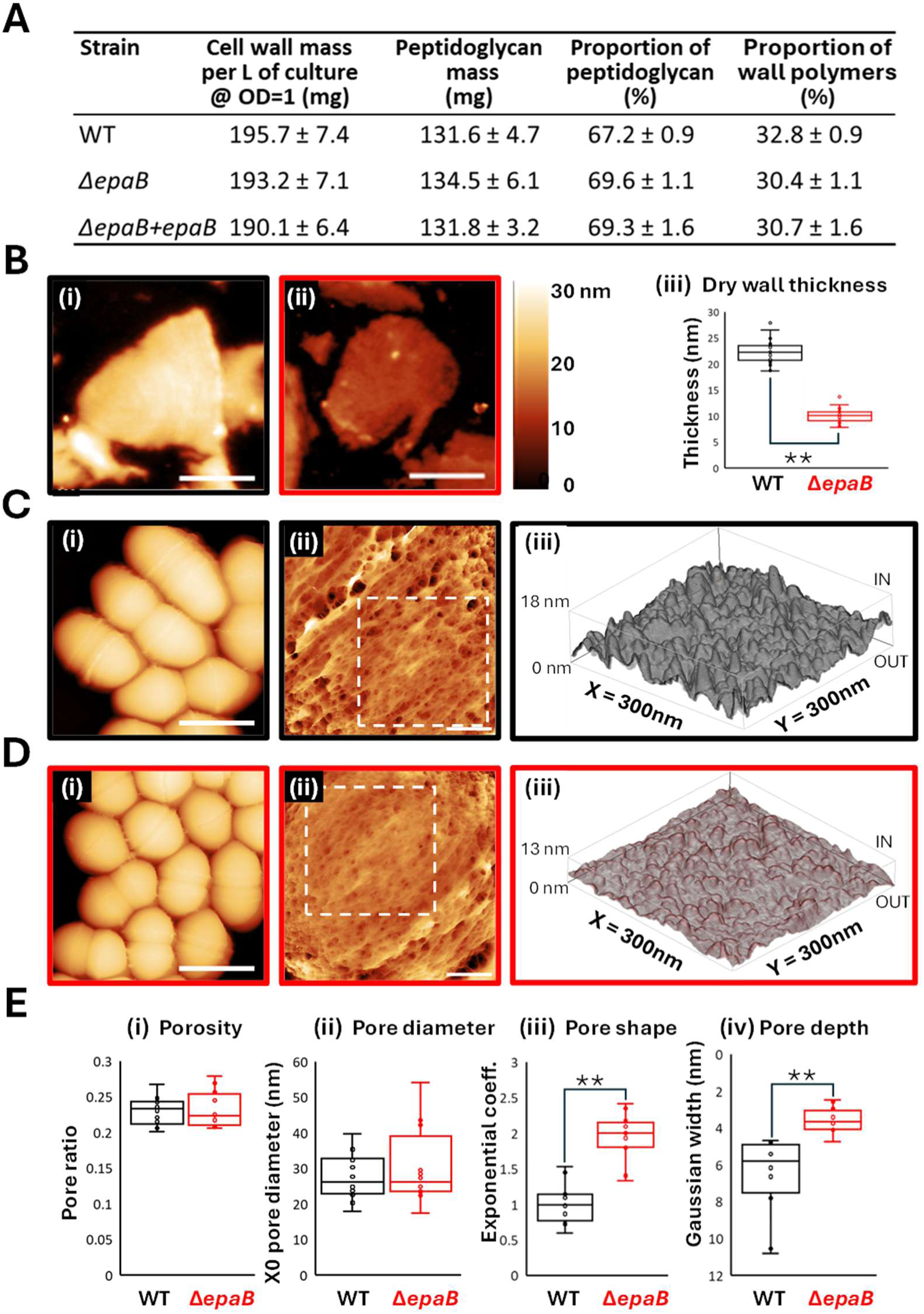
Comparing cell wall ultrastructure in *E. faecalis* OG1RF wild-type and *E. faecalis* OG1RF Δ*epaB* using atomic force microscopy. (**A**) Comparison of the proportion of peptidoglycan and cell wall polymers of wild type (WT), Δ*epaB*, and complemented strain (Δ*epaB + epaB*), with all values normalised to 1L of culture at OD600=1. No statistical differences between cell wall masses, and proportions of peptidoglycan cell wall polymers, were found across any pairwise comparison, based on a one-way ANOVA. (**B**) Dry cell wall fragments (peptidoglycan and polymers) of WT (i) and Δ*epaB* (ii) normalised against a 0-30 nm height scale, with thickness measured in air by AFM (iii) (WT = 22.35 ± 1.99 nm (n=42), Δ*epaB* = 10.01 ± 1.17 nm (n=50); *P* = 9.02E-44). (**C-D**) Whole cells imaged with AFM in liquid environment [scale bar=1 µm, data scale=700 nm];High resolution topography scans over the highest, flattest parts of the WT [C(ii)] and Δ*epaB* [D(ii)] cells (Scale bar = 100 nm, height scale = 30 nm); and 3-dimensional representations of the surface for the boxed areas shown in C (ii) and D (ii) to be analysed with AFMSlicer. (**E**). AFMSlicer analyses of: **E (i)** Porosity (WT = 0.2305 ± 0.0199, Δ*epaB* = 0.2313 ± 0.0251; *P* = 0.9362 – no significant difference); **E (ii)** X0 Pore diameter (WT = 27.62 ± 6.57 nm, Δ*epaB* = 30.08 ± 10.82 nm; *P* = 0.5093 – no significant difference); **E (iii)** Pore Shape (WT = 1.021 ± 0.279 – more cylindrical, Δ*epaB* = 1.953 ± 0.331 – more conical; *P* = 1.85E-07); **E (iv)** Pore Depth (WT = 6.541 ± 2.148, Δ*epaB* = 3.576 ± 0.672; *P* =4.42E-04). In **E**, n = 12 for WT and Δ*epaB*.

### The rhamnose backbone is essential for virulence and underpins phagocyte evasion

We used the zebrafish model of infection to explore the role of the EPA rhamnan chain in pathogenesis. The Δ*epaB* mutant showed a significant decrease in virulence as compared to the parental OG1RF strain and virulence was restored by complementation (Fig. 7A). Infection of transgenic zebrafish embryos expressing mCherry in phagocytes showed that the Δ*epaB* mutant was more readily taken up by phagocytes as compared to the parental OG1RF strain or its complemented derivative (Fig. 7B). Quantification of uptake was carried out by measuring the fluorescence intensity ratio between the signal associated with bacteria (expressing Green Fluorescent Protein) and macrophages (expressing mCherry). It confirmed a clear defect in innate immune evasion in the *epaB* mutant, which was more readily uptaken by macrophages as compared to the WT and complemented derivative (Fig. 7C and Fig. S4).

**Fig 7.**
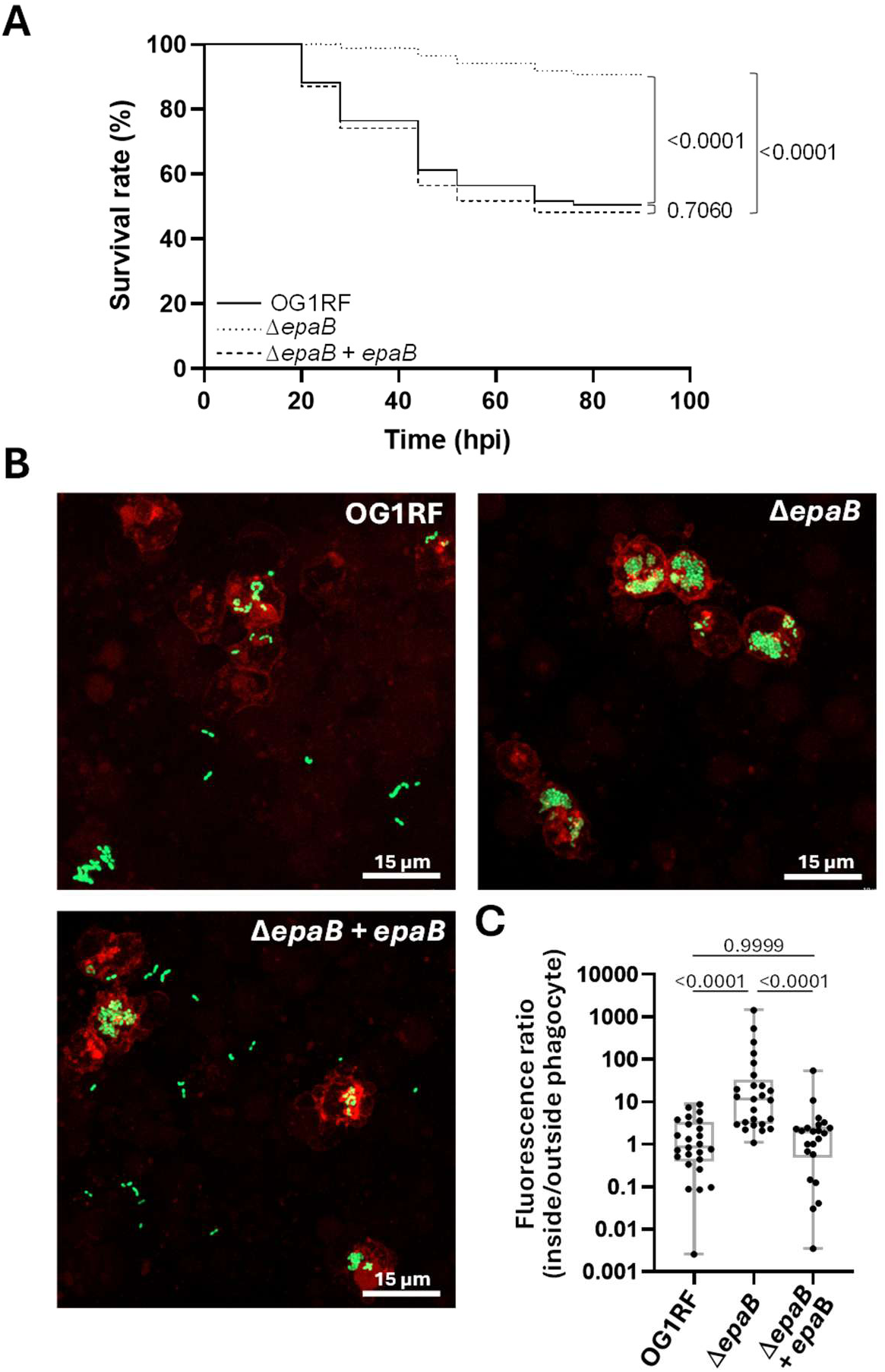
Virulence and host-pathogen interactions are impaired in the in the Δ*epaB* mutant. (**A**) Survival of zebrafish larvae following injection with *c.a.* 1500 CFUs of strains OG1RF, Δ*epaB* and complemented derivative (Δ*epaB* + *epaB*). Data shown combines three independent experiments, each corresponding to the injection of ≥25 (individual repeats are shown in supp Fig. 4). (**B**) Uptake of *E. faecalis* OG1RF (top panel), Δ*epaB* (right panel) and complemented derivative (bottom panel) by zebrafish macrophages. Embryos of the *Tg*(*mpeg:mCherry-F*) transgenic line were infected with c.a. 1500 *E. faecalis* cells expressing GFP and fixed in 4% (w/v) paraformaldehyde 1.5 hours post infection. Fluorescent bacteria and phagocytes were imaged by scanning confocal microscopy. Representative images of phagocytes following infection with wild type OG1RF, Δ*epaB* or complemented Δ*epaB* mutant (Δ*epaB* + *epaB*) are shown. Macrophages appear in red, GFP-producing bacteria in green. The scale bar is 10 µm. The area of GFP fluorescence signal outside and inside macrophages was measured and the ratio of GFP fluorescence area inside to outside phagocytes was used to quantify bacterial uptake. (**C**) Pairwise comparisons of fluorescence ratios was compared using an unpaired non-parametric Dunn’s multiple comparison test. Phagocytosis was significantly higher for the Δ*epaB* mutant when compared to the parental OG1RF (P<0.0001) or the complemented strain (P<0.0001). No difference in uptake was found between the parental strain OG1RF and complemented mutant (P=0.999).

## DISCUSSION

*epaB* is the second gene of the locus proposed to encode the rhamnose backbone (Fig. 1A). *E. faecalis* mutants carrying either a transposon insertion or an in-frame deletion in the *epaB* gene have been reported in the literature (17, 24). Because earlier studies describing the *epaB* transposon mutant did not systematically include complementation experiments, and because the structure of the EPA_B polysaccharide remains unresolved, we performed a comprehensive characterisation of this mutant. We have now elucidated the structure of the EPA polymer in the Δ*epaB* mutant and demonstrated that it produces a surface EPA polymer that contains only the decorations directly linked to the MurNAc residue of the peptidoglycan. The absence of the rhamnan chain in EPA is associated with alterations in cell wall architecture. Despite the presence of decorations known to be essential for virulence and for bacteriophage infection, our findings indicate that the rhamnan chain itself is also required for the full biological activity of EPA.

Composition analysis of OGR1F and its *epaB* derivative revealed the lack of rhamnose in the mutant cell walls and the presence of mannose (24). Whilst our results clearly demonstrate that EpaB is essential to produce the rhamnan chain, we could not detect mannose in EPA. The presence of this sugar remains to be explained since OG1RF does not produce a capsule and no other polysaccharide has been reported in this strain. In both EPA_WT and EPA_B, the GalNAc residue at the non-reducing end of the decorations is linked to either a β-GlcNAc residue (at the end of the rhamnan chain) or an α-GlcNAc residue linked to peptidoglycan in EPA_B (Fig. 8).

**Fig 8.**
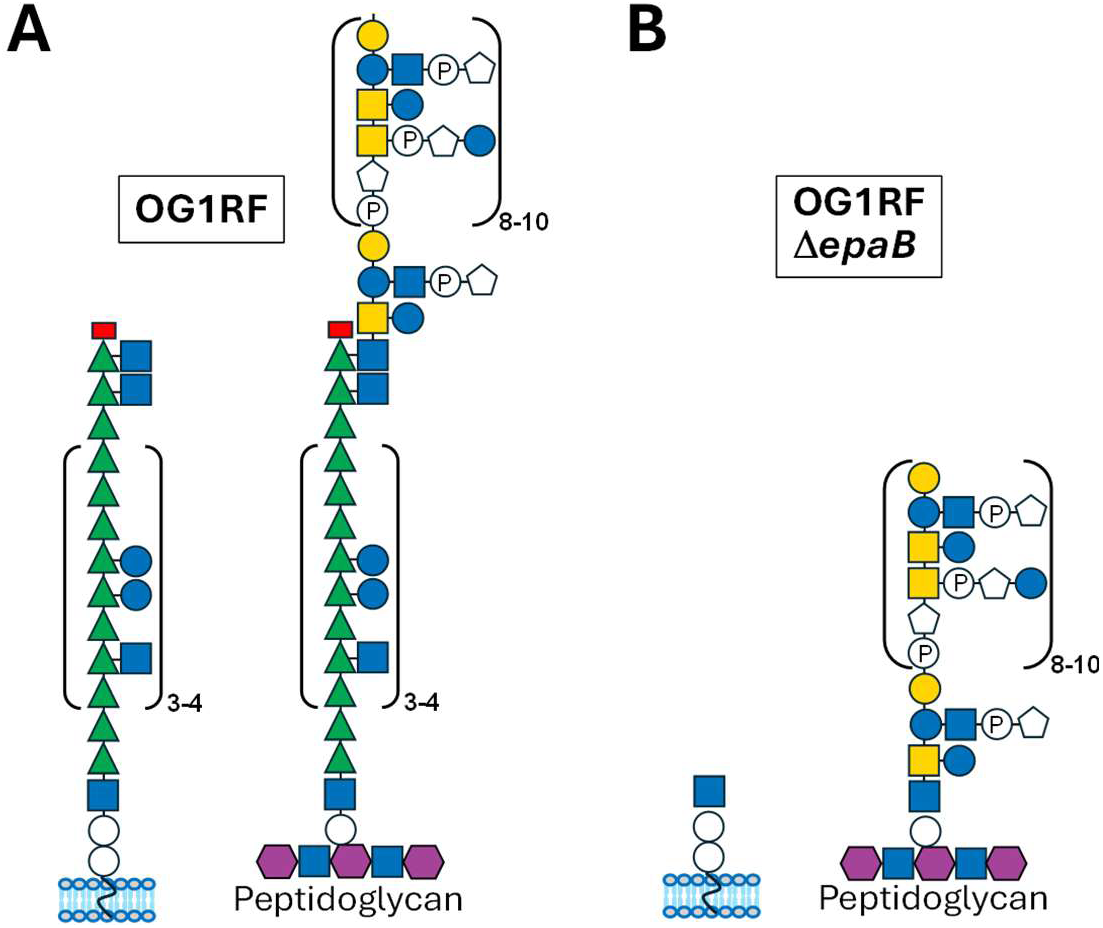
Biosynthesis of EPA_WT vs EPA_B. (**A**) In EPA_WT, the rhamnose backbone and the decoration region are synthesised as separate polymers, and the decorations are added to the terminal β-GlcNAc of the rhamnose backbone. (**B**) In EPA_B, in the absence of the rhamnose backbone, the α-GlcNAc of the peptidoglycan linker region, which canonically is an acceptor for the first residue of the rhamnose backbone, acts as an alternative acceptor for the decoration region. In both cases, an unidentified ligase covalently links the assembled polymer to the C6 position of the MurNAc residue in peptidoglycan.

This observation implies that the enzyme that links the GalNAc (residue M in Fig. 2) in EPA to the GlcNAc residue of an acceptor chain can accommodate distinct substrates. Likewise, the ligase that transfers the full-length EPA_WT or the EPA_B polysaccharide to the MurNAc residue of peptidoglycan must be able to accommodate these two different substrates. Five LytR-Cps2A-Psr (LCP) homologs exist in OG1RF, however the enzyme(s) anchoring EPA to peptidoglycan have not been identified yet.

Thin-sectioning of the *epaB* mutant, revealed a pronounced morphological defect, with cells appearing rounder than the parental strain and exhibiting misplaced equatorial rings (24). However, the same authors could not detect differences in cell wall thickness, in contrast to the thinner cell wall observed using atomic force microscopy (AFM). The reason for this discrepancy remains unclear but may be attributed to fixation artefacts introduced during sample preparation and embedding for TEM versus the native state of samples in our AFM experiments. Despite the apparent reduction in cell wall thickness in the *epaB* mutant observed by AFM, EPA still accounts for *c.a.* 30% of the cell wall dry weight, a proportion comparable to that of the wild-type strain (Fig. 6A). This finding suggests a compensatory mechanism in which the absence of the rhamnan chain is offset by enhanced decoration levels, therefore changing the cell wall organisation and affecting the dry thickness measured by AFM. Based on the quantification of NMR signals, truncated EPA_B molecules present a similar degree of polymerisation to EPA_WT. Δ*epaB* mutants therefore do not compensate for the lack of a rhamnan chain with longer decoration chains. Interestingly, the cell wall architecture of the Δ*epaB* mutant is altered, causing a round cell shape phenotype as seen by high-resolution AFM in liquid environment of non-fixed cells whole (Fig. 6 C-D). This dramatic change in cell shape is in agreement with the phenotype observed previously with TEM (24). Recent improvements in AFM imaging methodology (23), has allowed for the nanometric architecture of the cell wall *E. faecalis* to be revealed at unprecedented resolution in this study. The glycan strands from the peptidoglycan create a fibrous mesh with pores in between. By quantitatively analysing (https://zenodo.org/records/13133978) several high-resolution images (n=12 for each strain), we showed that the *epaB* mutant has indeed undergone a reorganisation of its cell wall nanometric architecture, with significantly shallower and more conical cell wall pores (Fig. 6E). Although monitoring the surface display of EPA decorations is challenging, the observed resistance to bacteriophages that rely on specific structural motifs of these decorations to infect *E. faecalis* OG1RF suggests that the absence of rhamnan alters EPA surface presentation. This agrees with the AFM results suggesting a re-organisation of the cell wall components. Together with previous studies demonstrating that EPA decorations are essential for bacteriophage infection (10, 24–27), these findings indicate that both components of the EPA polymer are required for efficient bacteriophage-induced killing.

Several studies have demonstrated that intact EPA decorations are essential for host colonisation (28), biofilm formation (7), antimicrobial resistance (9), and pathogenesis in various experimental infection models (9, 24). Mutations in the conserved *epa* locus, which encodes the rhamnan backbone, have also been associated with reduced virulence; however, the lack of detailed structural data describing the effects of these mutations has led to conflicting interpretations, with some studies claiming that *epaB* mutants no longer produced EPA (16–18). We show here that *epaB* mutants produce intact decorations but remain avirulent and are unable to evade innate immune evasion in the zebrafish model of infection. *epaB* mutants display a slight resistance to SDS and lysozyme but no resistance to polymyxin B or sodium cholate like EPA decoration mutants do (Fig. S5 and (9)). Analysis of the *epa* locus led us to hypothesise that the rhamnan backbone and its decorations are produced independently. We proposed that the rhamnan chain is translocated across the cytoplasmic membrane as a fully assembled polymer via a Wzx/Wzy-like pathway, whereas the decorations are translocated via an ABC transporter and polymerised at the cell surface (11).

The existence of two independent polysaccharide chains encoded by adjacent genetic loci is conserved in *Lactococcus lactis*, where a rhamnan polymer is produced alongside another polysaccharide termed the pellicle (29, 30) Although no structure has yet been determined for the full-length, intact polysaccharide produced by *L. lactis*, it has been proposed that the rhamnan and pellicle polymers are covalently linked (29). Surprisingly, *E. faecalis epaB* ortholog *rgpB* in *L. lactis* could not be deleted and depletion experiments revealed a profound impact on cell viability (30). Further investigations are therefore needed to compare the structure/function of the rhamnopolysaccharides in these related lactic bacteria.

## MATERIALS AND METHODS

### Bacterial strains and growth conditions

Bacterial strains used in this study are described in Table S4. *E. faecalis* was grown in Brain Heart infusion media at 37°C without agitation. Tetracycline was added at 5 μg/mL to maintain plasmid pMV158-GFP.

### Concanavalin A staining

Cultures were inoculated in LB media with 100 µg/ml carbenicillin and grown at 37°C under agitation (200 rpm) until an OD600 of 0.6 was reached. Expression of recombinant proteins was then induced with 0.5% (w/v) arabinose and cultures were incubated for 16 h at 28°C. The next day, 0.1 ml of cells diluted to an OD600 of 0.3 were centrifuged at 3,300 x g for 3 mins at 4°C. Pellets were washed twice with 1 ml of filtered and autoclaved PBS (137 mM NaCl, 2.7 mM KCl, 10 mM Na2HPO4, 1.8 mM KH2PO4) and cells were incubated for 30 minutes in the presence of 5 µg/ml Concanavalin A conjugated to Alexa Fluor 633 (Thermo Fisher Scientific) to label the *O*-antigens. Cells were washed twice and fixed in 1 ml of 2% (w/v) paraformaldehyde for 15 minutes at 4°C. After two washes with 1 ml of filtered and autoclaved PBS supplemented with 0.1% Triton X-100, cells were resuspended in 1 ml of filtered PBS supplemented with 1% (w/v) BSA, diluted to an OD600 of 0.1 in the same buffer and pipetted in triplicate into a flat bottom 96 well plate for flow cytometry analysis.

### Flow cytometry analysis

Stained and fixed cells were analysed using a NovoCyte cytometer (Agilent). Cell fluorescence was detected using the forward scatter laser (640 nm) and a 675/30 nm filter. The data generated was analysed using FlowJo **(V10.10)** software.

### Cell wall, peptidoglycan and EPA purification

Cell walls were isolated from 1.8 L of exponentially growing cultures by boiling in 4% (w/v) SDS for 30 minutes. The material was washed six times with Milli-Q water and incubated with DNase and RNase (50 µg/mL each) in 20 mM Tris-HCl (pH 7.5), 150 mM NaCl, and 5 mM MgCl₂ for 2 hours at 37 °C. Pronase was then added to 1 mg/mL and incubated for 3 hours at 55 °C. Enzymes were inactivated by addition of SDS to 1% (w/v) followed by boiling for 20 minutes. Cell walls were washed six times with Milli-Q water and freeze-dried. Peptidoglycan-anchored polymers were removed by treatment with 1 M HCl at 37 °C for 5 hours. EPA was purified as previously described by size-exclusion and anion exchange chromatography following cell wall digestion with mutanolysin (11).

### Solution-state NMR experiments

NMR spectra were recorded on a 600 MHz NEO using a 5-mm-diameter TCI cryo-probe (^1^H and ^13^C) and a 500 MHz Bruker Avance II using a 5-mm-diameter BBO probe (^1^H, ^13^C, and ^31^P). All experiments were performed at 298 K in D2O. Samples were supplemented with 0.01 % (v/v) acetone as a reference to calibrate ^1^H and ^13^C chemical shifts, which are reported as parts per million (δH 2.225 ppm and δC 31.55 ppm). ^1^H-^1^H COSY experiments used 2D homonuclear shift correlation using gradient pulses for selection and presaturation during the relaxation delay. ^1^H-^1^H TOCSY experiments were recorded using mixing times of 20, 40, 60, 80, 100, and 120 ms across multiple experiments. ^1^H-^1^H NOESY NMR experiments were recorded using mixing times of 50 ms. A phase-sensitive multiplicity-edited sequence was used to record the ^1^H-^13^C HSQC which used echo/anti-echo-TPPI gradient selections with decoupling during acquisition and trim pulses in the Inept transfer. The ^1^H-^13^C HSQC-TOCSY NMR experiments were acquired using MLEV17 for homonuclear Hartmann-Hahn mixing and echo/anti-echo-TPPI gradient selections with decoupling during acquisition and trim pulses in the Inept transfer, and a mixing time of 100 ms. The ^1^H-^13^C HMBC NMR experiments were optimised on long-range couplings *^n^J*CH of 8 Hz without multiplicity selection or decoupling during acquisition. The ^1^H-^31^P HSQC-TOCSY NMR experiments were phase sensitive and acquired using MLEV17 for homonuclear Hartmann-Hahn mixing with decoupling during acquisition and trim pulses in the Inept transfer and a mixing time of 30 ms. The ^1^H-^31^P HSQC-TOCSY NMR experiment was phase-sensitive and acquired using MLEV17 for homonuclear Hartmann-Hahn mixing with decoupling during acquisition with a mixing time of 40 ms. Spectra were processed on Topspin 3.2 and analysed on Topspin 4.4.1.

### Solid-state (HR-MAS) NMR

A culture corresponding to 100 mL of cells in the exponential phase was spun and heat-killed at 60 °C for 20 minutes. Cells were then washed twice in D2O, and freeze-dried. Dried cells were re-suspended into 100 μL of D2O containing 0.01% (v/v) acetone as an internal standard for chemical shifts (δ^1^H 2.225 and δ^13^C 31.55) and centrifuged at 3,000 rpm to be packed into a 4-mm ZrO2 rotor (CortecNet, Paris, France) (31). HR-MAS NMR was performed using an 18.8T Avance NEO spectrometer where ^1^H and ^13^C resonated at 800.12 and 200.3 MHz respectively. The set of pulse programs used was extracted from the Bruker pulse program library where pulses (both hard and soft pulses and their powers) and delays were optimized for each experiment. ^1^H-^13^C HSQC spectra were recorded with an inept sequence to distinguish secondary carbons and carbons bearing primary alcohols from other carbons. The 18.8 T was equipped with a 4mm D/^1^H/^13^C/^31^P HR-MAS probe head where the rotor was spun at 8 kHz during acquisition to eliminate the anisotropy effect of jelly state of bacterial cells. All spectra were recorded at 300 K, and the rotor spinning rate was 8 kHz. For ^1^H-^13^C HSQC experiments, the spectral widths were 12,820 Hz (^1^H) with 1,024 points for the FID resolution and 29,994 Hz (^13^C) with 400 points for FID resolution during 400 scans, giving 12.5 Hz/pt and 75.0 Hz/pt, respectively.

### Sample preparation for AFM

Cells were grown to OD600=0.7, heated at 100°C for 30 minutes, washed in phosphate saline buffer (PBS) three times and resuspended at a final OD600 of 20. For sacculi preparation, boiled cells stocks were diluted 20-fold in PBS to a final volume of 1 mL then transferred to 2 mL Lysing matrix tubes with 0.1 mm Silica beads. Cells were mechanically broken by 6 cycles of bead-beating at maximal speed for 30 seconds in MP Biomedical^TM^’s FastPrep^®^-24 5G Bead beating system. Silica beads were removed by a series of centrifugations at 100 x *g* for 1 minute. Bead-free cell lysates were centrifuged at 17,000 x g, resuspended in 5% (w/v) Sodium dodecyl sulphate (SDS) and boiled at 100°C on a heat block for 30 minutes. After 3 washes in MilliQ water, cells were boiled in SDS again for a further 15 minutes. Lysates were washed 5 times with HPLC-grade water and resuspended in 900 μL of 20mM Tris-HCl (pH 7.5) and 100 μL Pronase then incubated at 60°C on a heat block for 90 minutes. Cell wall sacculi resulting from this treatment were washed with HPLC-grade water and resuspended to an OD600 of 5.0. 8mm mica discs were adhered to 12mm metal pucks for AFM using Flexbar’s *Reprorubber Thin-pour Green Glue*. Once adhered, individual mica layers were cleaved off with *Scotch Magic^TM^ Tape* until a uniform flat surface remained. 100 µL of 0.1% Poly-L-Lysine (PLL) were added on top of the cleaved mica and incubated at room temperature for 20 minutes. PLL was removed and the mica was rinsed with 100 µL of HPLC-grade water before being blown dry with sterile air flow (Prepared mica). 20 µL of boiled cell stock were added to prepared mica and incubated at room temperature for 5 minutes. This was removed by rinsing with 100 µL of HPLC-grade water then blown dry with sterile air. Samples were left dry for at least 24 hours before imaging.

Sacculi stock was diluted 5-fold in PBS to a final volume of 100 µL and using the smallest probe possible from *Soniprep 150 plus, MSE*, the diluted stock was subject to two cycles of 30 seconds at 2 mA. 20 µL of diluted, sonicated sacculi stock was incubated onto prepared mica for 5 minutes at room temperature. This was removed by rinsing with 50 µL of HPLC-grade water then blown dry with sterile air. Samples were left dry for at least 24 hours before imaging.

### AFM Image Acquisition

Imaging of boiled cells in liquid was performed on a *Bruker’s Dimension Fastscan* instrument in a buffer containing 150 mM NaCl, 10 mM Tris (pH 7) and 50 mM MgCl2 and *Bruker Fastscan-D* probes with a nominal spring constant of 0.25 Nm^−1^ and resonant frequency of 110 kHz. Images were constructed using *ScanAsyst^TM^ Peakforce (PF) mapping* with a typical PF amplitude of 100-200 nm, scanning frequency of 2 kHz and set point between 1-2 nN. Sacculi were imaged in air using a *Bruker’s Veeco Multimode 8* instrument under ambient conditions and *NuNano Scout 350-RAI* probes with spring constants of 42 Nm^−1^. Cantilevers were tuned just below resonance with a free amplitude of ∼25nm, then imaged under a set-point amplitude of 50-70% of the free amplitude. This corresponds to tip-sample imaging forces below 500 pN.

### AFM data analysis

High-resolution AFM images were set to greyscale, cropped to squares using the open-source *Gwyddion* software (32) to isolate qualitatively uniform areas of peptidoglycan mesh. Crops were flattened with polynomial (2^nd^ degree) row alignment and subject to a final removal of horizontal scars. The images were then exported as .tiff 16-bit files, the option of “none” was chosen for the colour and scale bars (no frame was added). These slightly flattened images were then imputed on the AFMSlicer (https://zenodo.org/records/13133978) with macros 1 in FIJI, slicing the images in 256 slices along the Z axis. The stack of all the slices was given their corresponding XYZ dimensions, extracted previously from Gwyddion > dimensions tool. This stack was used by AFMSlicer macro 2 in FIJI to systematically and automatically analyse each individual pore in the image and provide the four pore characteristics presented in Figure 6 (pore diameter, porosity, pore depth and conicality).

Cell wall thickness was measured using the *1D statistical functions* tool in *Gwyddion* as previously described (33).

### Phage infection assays

Efficiency of plating was determined using the agar overlay method, as previously described (Al-Zubidi et al., 2019). *E. faecalis* indicator strains were grown to exponential phase (OD600≈0.5) in BHI supplemented with 5 mM MgSO4 and 5 mM CaCl2 and mixed various dilutions of phage stocks adjusted at 10^8^ Plaque Forming Units (PFU)/mL. After 10 minutes at room temperature, the phage/bacteria mixture was added to 4 mL of BHI-top agar (0.6 % w/v) and immediately spread on a BHI-agar plate. The number of PFU/mL measured with OG1RF as an indicator strain was used to define 100 % infection.

### Zebrafish survival experiments

Zebrafish embryos were obtained by the natural spawning of adult zebrafish (line AB/TL), which were housed in a continuous recirculating closed-system aquarium with a light/dark cycle of 14/10 hours at 28 °C. Dechorionated embryos at 30 hours post fertilization (hpf) were anaesthetized and systemically injected with ca. 1,500 *E. faecalis* cells as previously described (13). The number of injected bacterial cells was checked before and after each series of injections with a given strain. The infected embryos were monitored at regular intervals until 90 hours post infection (hpi). At least 25 embryos per group were used in each experiment.

### Imaging of infected larvae by confocal microscopy and quantification of bacterial uptake by phagocytes

Larvae from the *Tg*(*mpeg:mCherry-F)* (34) transgenic line were fixed in 4% (w/v) paraformaldehyde at 1.5 hpi. Fixed larvae washed with PBS were then immersed in 1% (w/v) low-melting-point agarose solution in E3 medium and mounted flat on a glass-bottomed dish. Images were acquired with a Zeiss LSM900 Airyscan 2 confocal laser scanning microscope using the C-Apochromat 40x/NA 1.2 water objective. Maximum projections were used for representative images. No non-linear normalisation was performed. Bacterial phagocytosis was quantified as previously described (35). Briefly, all bacterial clusters were identified based on their GFP fluorescence. Next, the fluorescence intensities of mCherry-labelled macrophages surrounding the bacteria (2 μm radius) were analysed using a custom ImageJ script called Fish Analysis v5 (http://sites.imagej.net/Willemsejj/). The phagocytosed bacteria had a high intensity of mCherry fluorescence (inside the macrophages) in the surrounding area, and the cutoff of 2 times the background level was used to distinguish the phagocytosed from non-phagocytosed bacteria. The area of phagocytosed bacteria was compared to the area of non-phagocytosed bacteria and their ratio was calculated.

### Statistical analyses

Statistical analyses were performed using GraphPad Prism. Comparison between conA labelled samples and the proportions of cell wall peptidoglycan and polymers were made using an ordinary one-way ANOVA. Survival experiments were evaluated using the Kaplan-Meier method. Comparisons between curves were made using the Log Rank (Mantel-Cox) test. For macrophage uptake experiments, fluorescence intensity ratios were compared using an unpaired non-parametric Dunn’s multiple comparison test. The statistical analysis performed on the AFM data (thickness and nanometric architecture analysis) were unpaired student *t*-tests with Welch’s correction.

## ACKNOWLEDGEMENTS

LK, JLD and RES were funded by the White Rose Doctoral Training Programme (BBSRC grants BB/T007222/1, BB/T007222/1 and BB/M011151/1, respectively). KC and JSN were supported by PhD studentships from the DiMeN Doctoral Training Programme (Medical Research Council grants MR/W006944/1 and MR/N013840/1, respectively). NN is supported by a studentship from the University of Sheffield Faculty of Science. LPL was funded by the Wellcome Trust (Early Career Award 227581/Z/23/Z) and the Royal Society (Research Grant RG\R1\251553). TL was supported by a BBSRC EASTBIO studentship; HCD was additionally supported by a Wellcome Trust Career Development Award (225350/Z/22/Z). BJM and TKP were supported by the National Science Centre of Poland within Sonata Bis 9 project (Grant number UMO-2019/34/E/NZ6/00137). The authors would like to thank Dr. B. Doumer and Dr. J. Trebosc for their help on the NMR facility (800 MHz spectrometer) of the Advanced Characterization Platform of the Chevreul Institute. An upgrade to the 600 MHz NMR spectrometer at the University of Sheffield Biomolecular NMR Facility was funded by the Biotechnology and Biological Sciences Research Council (BB/R000727/1). Prof. Dirk Haller (Technical University of Münich) is acknowledged for the kind gift of the Δ*epaB* mutant and its complemented derivative.

## SUPPLEMENTAL MATERIAL

This article has 5 supplemental figures and 4 supplemental tables.

## DATA AVAILABILITY

All the bacteriophages, bacterial strains and plasmids described in Table S4 are available upon request to the authors.

## ETHICS APPROVAL

The Jagiellonian University Zebrafish Core Facility (ZCF) is a licensed breeding and research facility (District Veterinary Inspectorate in Krakow registry; Ministry of Science and Higher Education record no. 022 and 0057). All larval zebrafish experiments were conducted in accordance with the European Community Council Directive 2010/63/EU for the Care and Use of Laboratory Animals of Sept. 22, 2010 (Chapter 1, Article 1 no.3) and Poland’s National Journal of Law act of Jan. 15, 2015, for Protection of animals used for scientific or educational purposes (Chapter 1, Article 2 no.1). All experiments with zebrafish were done on larvae up to 5 days post fertilization, which have not yet reached the free feeding stage, and were performed in compliance with ARRIVE guidelines.

## SUPPLEMENTAL MATERIAL

**Supplemental Figure S1.**
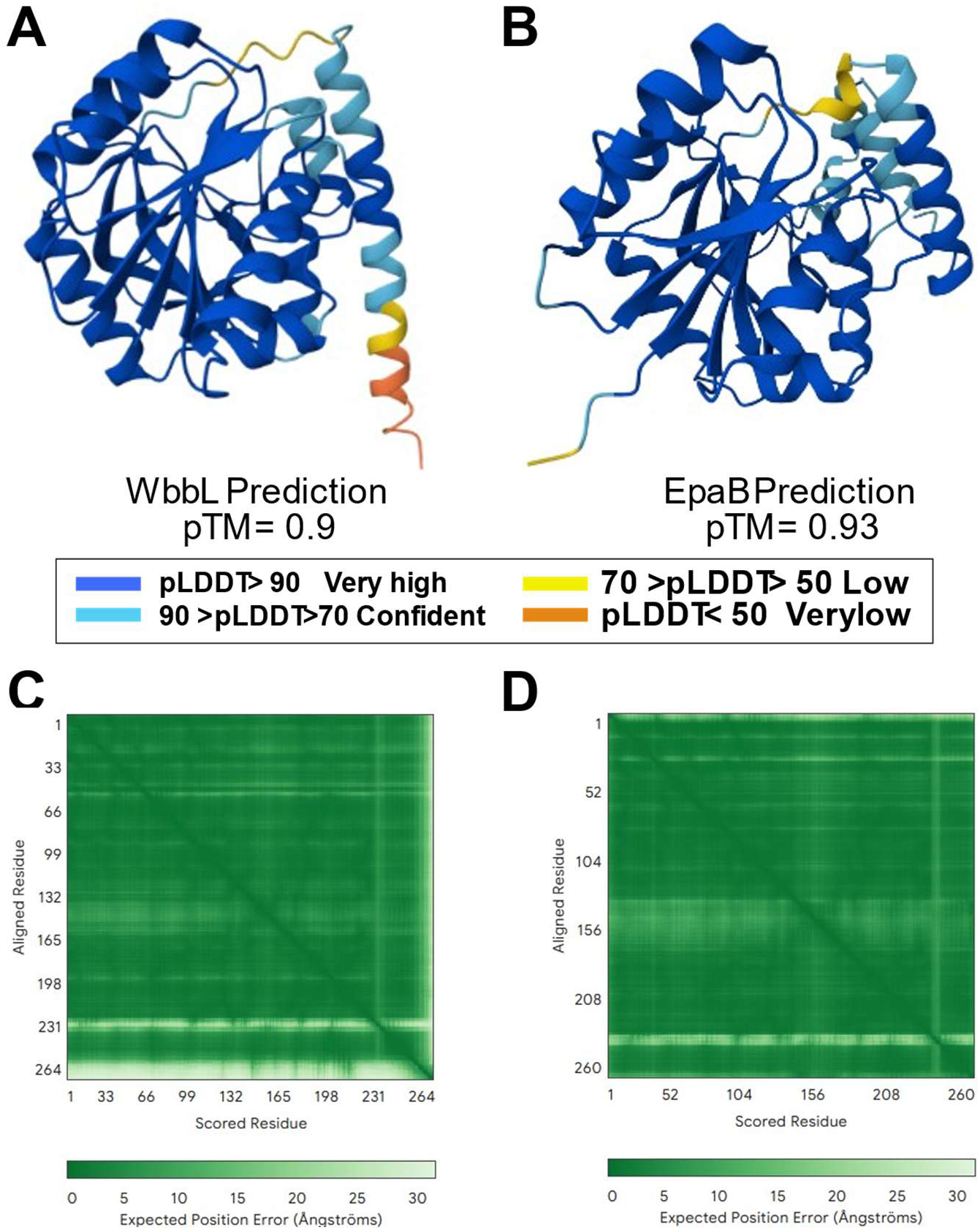
AlphaFold predictions of *E. coli* WbbL and *E. faecalis* EpaB. **A, B**, structural models reveal very high confidence predicted Local Distance Difference Test values (pLDDT). **C,D**, Expected Position Error values across the entire protein sequences.

**Supplemental Figure S2.**
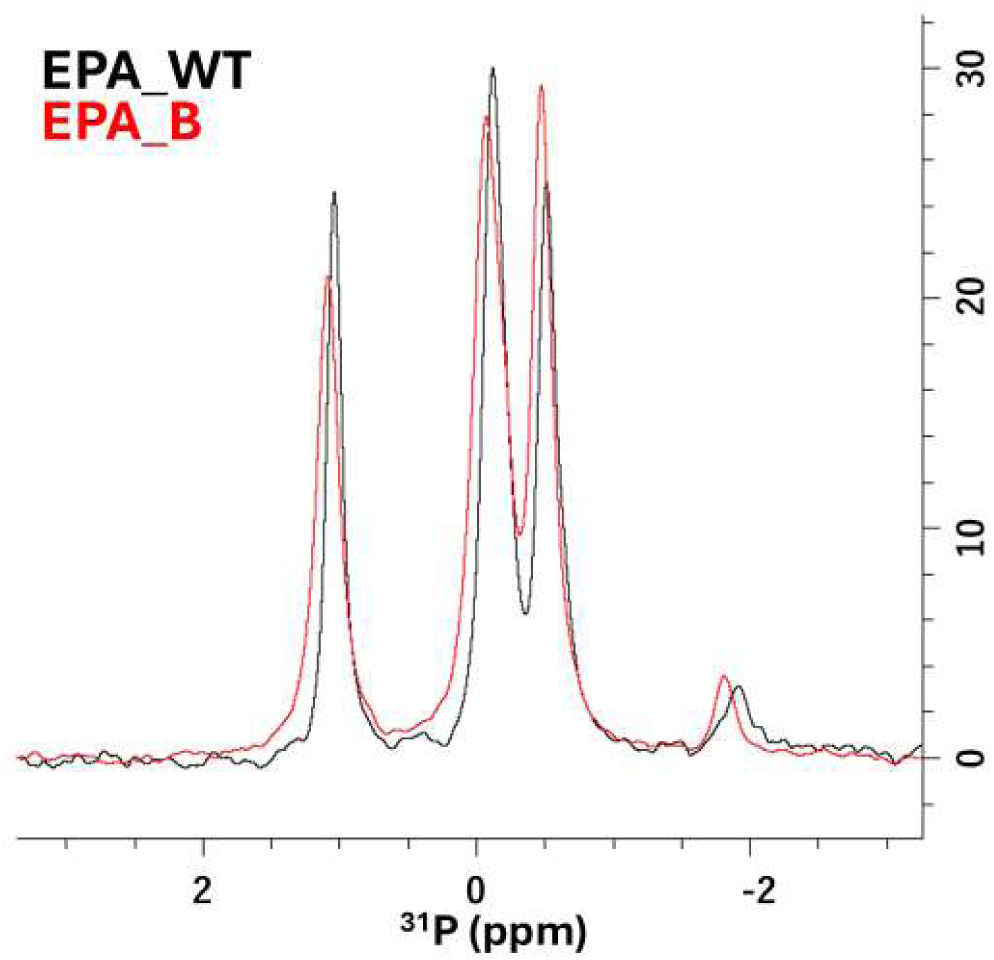
1D ^31^P spectrum of EPA_WT (black) and EPA_B (red). Phosphorus signals in both polysaccharides correspond to the phosphorus signals previously described, suggesting that they contain identical decorations.

**Supplemental Figure S3.**
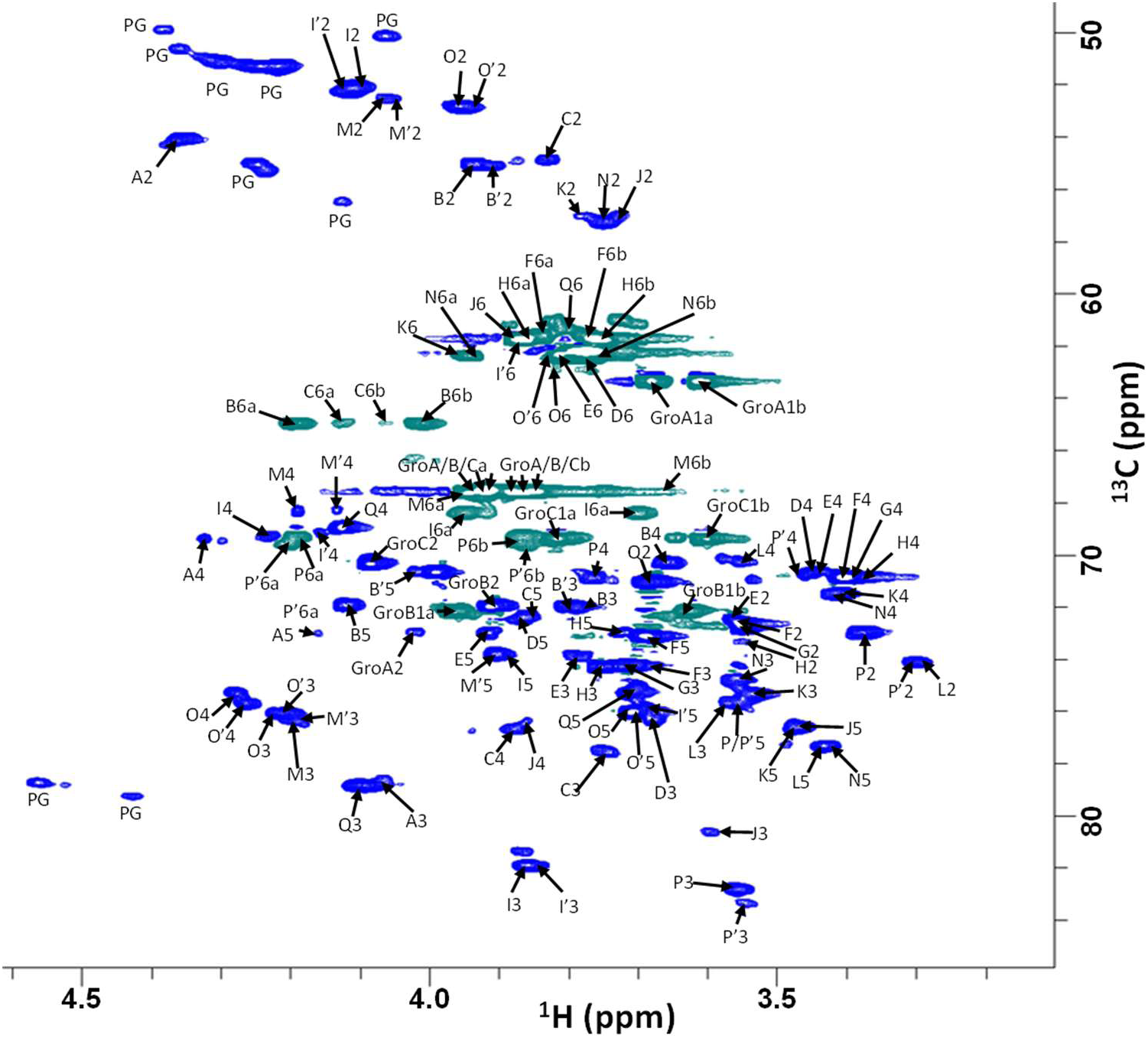
Assignment of the ^1^H-^13^C HSQC EPA_B spectrum.

**Supplemental Figure S4.**
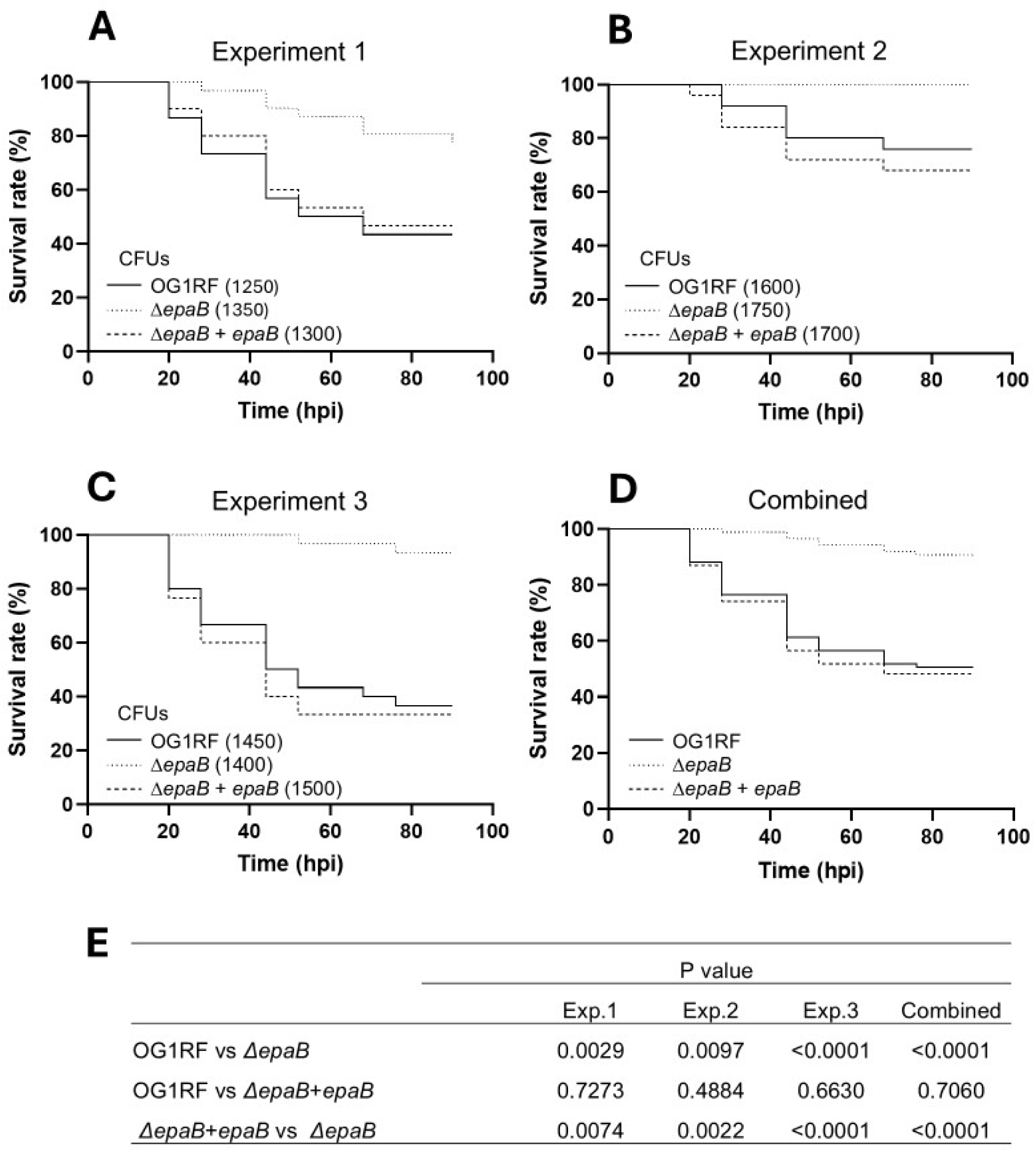
Survival ratio of zebrafish larvae infected with *E. faecalis* OG1RF, Δ*epaB* and Δ*epaB* complemented strains. Larvae were infected with *ca.* 1500 CFUs of parental (WT) OG1RF strain (solid line), Δ*epaB* (black dotted line) or Δ*epaB + epaB* (black dashed line). Survival was monitored between 20 to 90 hours post infection (hpi) at 28°C using ≥25 larvae per strain per experiment. Three independent experiments (**A**, **B** and **C**) and combined results (**D**) are shown. (**E**) P values of pairwise comparison.

**Supplemental Figure S5.**
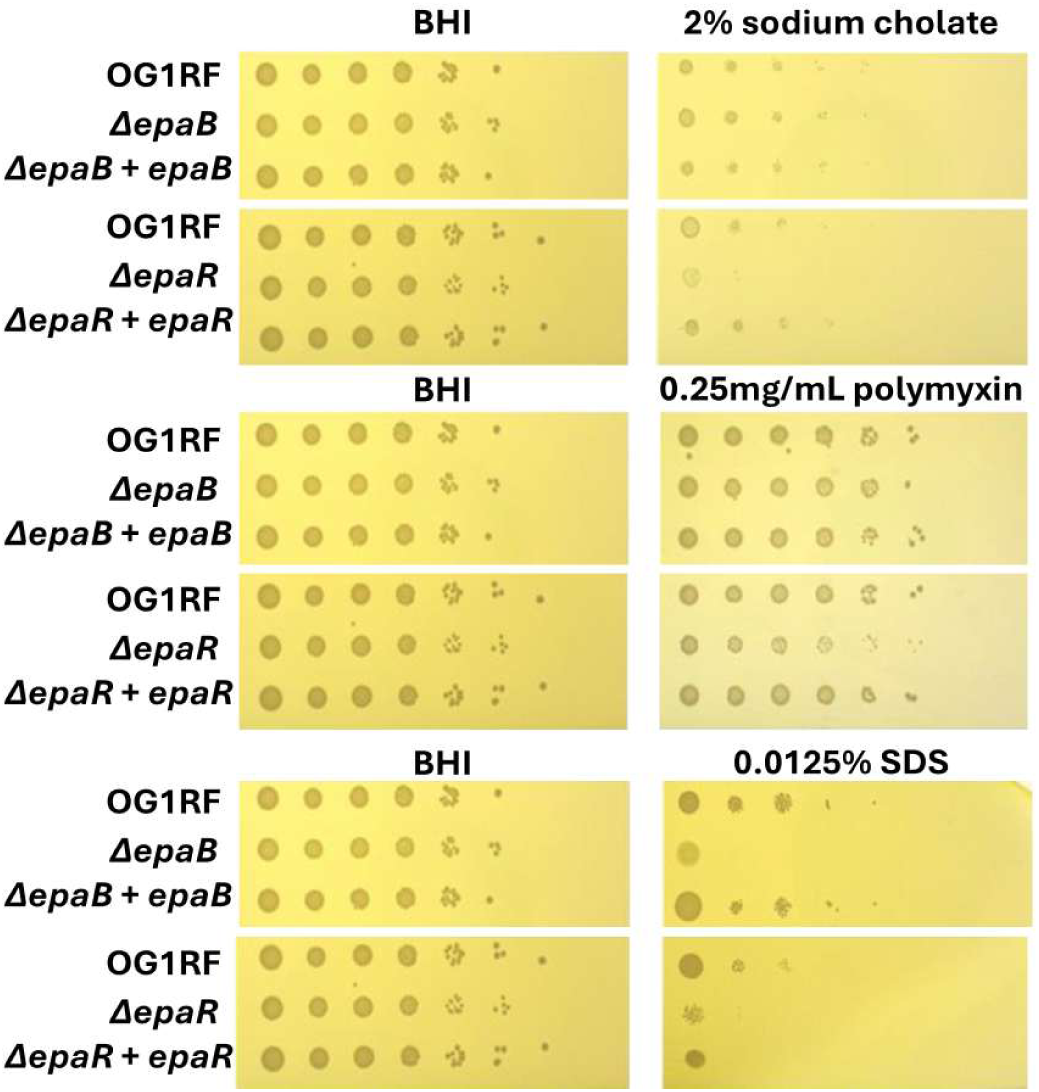
Antimicrobial susceptibility of the OG1RF, Δ*epaB* and complemented derivative. Serial dilutions of an overnight cultures were adjusted to an OD600nm at 1.0 and serially diluted (10-fold) in phosphate buffer saline. One microliter of each dilution was spotted on BHI (left panels) or BHI supplemented with various antimicrobials (2% (w/v) sodium cholate, 0.25 mg/mL polymyxin or 0.0125% (w/v) SDS) using a multipoint replicator. Pictures were taken 24h after incubation at 37°C.

**Supplemental Table 1:** All ^1^H, ^13^C and ^31^P NMR chemical shifts (ppm) of pyranose rings at 298 K of EPA purified from Δ*epaB* cells (EPA_B) determined through the analysis of ^1^H-^1^H-COSY, ^1^H-^1^H-TOCSY, ^1^H-^13^C-HSQC, ^1^H-^1^H NOESY, ^1^H-^13^C-HSQC-TOCSY, ^1^H-^31^P-HSQC and ^1^H-^31^P-HSQC-TOCSY experiments.

| Residue |  | H1<br>C1 | H2<br>C2 | H3<br>C3 | H4<br>C4 | H5<br>C5 | H6a<br>C6a | H6b<br>C6b | Nac<br>CH<sub>3</sub> | PO<sub>4</sub> |
| --- | --- | --- | --- | --- | --- | --- | --- | --- | --- | --- |
| → 3)α-GlcNAcp-(1→ | A | 5.52<br>95.75 | 4.37<br>54.10 | 4.05<br>78.78 | 4.33<br>69.50 | 4.17<br>72.90 | ND<br>ND | ND<br>ND | 2.06<br>23.72 | -1.83 |
| → 6)α-GlcNAcp-(1→ | B | 5.26<br>99.07 | 3.93<br>54.99 | 3.80<br>71.97 | 3.66<br>70.29 | 4.12<br>71.89 | 4.20<br>64.92 | 4.02<br>64.92 | 2.06<br>23.72 | 1.09 |
| α-GlcNAcp-(1→ | B' | 5.22<br>99.14 | 3.92<br>55.06 | 3.79<br>71.83 | 3.52<br>72.54 | 4.01<br>70.48 | ND<br>ND | ND<br>ND | 2.06<br>23.72 | 1.09 |
| → 4)-P-6-α-MurNAcp-OH | C | 5.24<br>91.43 | 3.83<br>54.83 | 3.74<br>77.61 | 3.87<br>76.59 | 3.86<br>72.32 | 4.13<br>64.95 | 4.07<br>64.95 | 2.06<br>23.72 | -1.83 |
| α-D-Glcp-(1→ | D | 5.18<br>96.64 | 3.71<br>73.00 | 3.68<br>76.42 | 3.44<br>70.88 | 3.86<br>72.50 | 3.75<br>61.90 | ND<br>ND |  |  |
| α-D-Glcp-(1→ | E | 5.10<br>97.17 | 3.56<br>72.51 | 3.79<br>73.80 | 3.45<br>70.63 | 3.91<br>73.10 | 3.82<br>62.36 | ND<br>ND |  |  |
| α-D-Glcp-(1→ | F | 4.97<br>99.64 | 3.56<br>72.52 | 3.71<br>74.20 | 3.40<br>70.65 | 3.68<br>73.20 | 3.85<br>61.64 | 3.75<br>61.64 |  |  |
| α-D-Glcp-(1→ | G | 4.96<br>99.45 | 3.55<br>72.65 | 3.72<br>74.38 | 3.39<br>71.04 | ND<br>ND | ND<br>ND | ND<br>ND |  |  |
| α-D-Glcp-(1→ | H | 4.94<br>99.51 | 3.55<br>72.83 | 3.74<br>74.33 | 3.38<br>70.84 | 3.70<br>73.02 | 3.85<br>61.65 | 3.75<br>61.65 |  |  |
| → 3,6)β-GalNAcp-(1→ | I | 4.71<br>103.61 | 4.11<br>52.03 | 3.86<br>81.73 | 4.23<br>69.29 | 3.40<br>73.92 | 3.95<br>68.37 | 3.82<br>68.37 | 2.06<br>23.72 |  |
| → 3)β-GalNAcp-(1→ | I' | 4.69<br>103.62 | 4.12<br>52.24 | 3.86<br>81.96 | 4.16<br>69.15 | 3.70<br>76.10 | 3.86<br>61.91 | ND<br>ND | 2.06<br>23.72 |  |
| → 4)β-MurNAcp-OH | J | 4.64<br>96.32 | 3.73<br>57.08 | 3.58<br>80.65 | 3.85<br>76.34 | 3.47<br>76.53 | 3.86<br>61.40 | ND<br>ND | 2.06<br>23.72 |  |
| β-GlcNAcp-(1→ | K | 4.63<br>101.62 | 3.78<br>57.05 | 3.54<br>75.14 | 3.40<br>71.51 | 3.48<br>76.80 | 3.94<br>62.39 | ND<br>ND | 2.06<br>23.72 |  |
| → 3)β-D-Glcp-(1→ | L | 4.59<br>105.52 | 3.31<br>74.04 | 3.57<br>75.66 | 3.56<br>70.21 | 3.45<br>77.26 | ND<br>ND | ND<br>ND |  |  |
| → 3,6)-β-D-GalNAcp-(1→ | M | 4.58<br>103.15 | 4.06<br>52.62 | 4.20<br>76.31 | 4.19<br>68.34 | 3.91<br>73.83 | 3.94<br>67.55 | 3.66<br>67.55 | 2.06<br>23.72 |  |
| → 3)β-D-GalNAcp-(1→ | M' | 4.56<br>103.15 | 4.059<br>52.62 | 4.18<br>76.15 | 4.13<br>68.26 | ND<br>ND | ND<br>ND | ND<br>ND | 2.06<br>23.72 |  |
| β-GlcNAcp-(1→ | N | 4.54<br>101.65 | 3.75<br>57.22 | 3.55<br>74.78 | 3.41<br>71.51 | 3.43<br>77.35 | 3.94<br>62.40 | 3.78<br>62.40 | 2.06<br>23.72 |  |
| → 3,6)β-GalNAcp-(1→ | O | 4.52<br>103.27 | 3.96<br>52.75 | 4.22<br>76.03 | 4.28<br>75.26 | 3.71<br>75.96 | 3.81<br>61.93 | ND<br>ND | 2.06<br>23.72 | -0.46 |
| → 3)β-GalNAcp-(1→ | O' | 4.51<br>103.27 | 3.95<br>52.90 | 4.20<br>76.21 | 4.27<br>75.72 | 3.71<br>76.05 | 3.81<br>61.82 | ND<br>ND | 2.06<br>23.72 | -0.46 |
| → 3,6)β-D-Glcp-(1→ | P | 4.52<br>105.92 | 3.38<br>72.91 | 3.55<br>82.83 | 3.37<br>70.94 | 3.56<br>75.61 | 4.19<br>69.29 | 3.86<br>69.29 | 2.06<br>23.72 |  |
| → 6)β-D-Glcp-(1→ | P' | 4.50<br>105.92 | 3.31<br>74.04 | 3.55<br>82.76 | 3.48<br>70.65 | 3.55<br>75.61 | 4.21<br>69.59 | 3.86<br>69.59 |  |  |
| 3→)β-D-Galp-(1→ | Q | 4.48<br>104.21 | 3.68<br>71.04 | 4.10<br>78.88 | 4.11<br>68.87 | 3.71<br>75.23 | 3.77<br>61.41 | ND<br>ND |  | -0.08 |

**Supplemental Table 2:** All ^1^H, ^13^C and ^31^P NMR chemical shifts (ppm) of glycerols at 298 K of EPA purified from Δ*epaB* cells (EPA_B) determined through the analysis of ^1^H-^13^C-HSQC, ^1^H-^13^C-HSQC-TOCSY, ^1^H-^31^P-HSQC and ^1^H-^31^P-HSQC-TOCSY experiments.

| Residue |  | H1a<br>C1 | H1b<br>C1 | H2<br>C2 | H3a<br>C3 | H3b<br>C3 | PO<sub>4</sub> |
| --- | --- | --- | --- | --- | --- | --- | --- |
| Glycerol-3-*P* | GroA | 3.68<br>63.33 | 3.61<br>63.33 | 4.02<br>72.92 | 3.94<br>69.59 | 3.88<br>69.59 | 1.09 |
| → 1)-Glycerol-3-*P* | GroB | 3.96<br>72.13 | 3.63<br>72.13 | 3.91,<br>71.94 | 3.93<br>67.52 | 3.87<br>67.52 | -0.08 |
| → 1)-Glycerol-3-*P* | GroC | 3.82<br>69.38 | 3.61<br>69.38 | 4.09<br>70.41 | 3.92<br>64.47 | 3.86<br>64.47 | -0.46 |

**Supplemental Table 3:** Residue connectivity. EPA purified from Δ*epaB* cells (EPA_B) determined through the analysis of ^1^H-^1^H NOESY and ^1^H-^13^C-HMBC experiments.

| Residue |  | NOE | HMBC |
| --- | --- | --- | --- |
| $\rightarrow 6)\alpha\text{-GlcNAcp-(1}\rightarrow$ | B | 5.25 → 3.55<br>B (H1) → P (H3) | 5.26 → 82.84<br>B (H1) → P (C3) |
| $\alpha\text{-D-Glcp-(1}\rightarrow$ | D | 5.18 → 3.82<br>D (H1) → GroC (H1a) | 5.18 → 69.31<br>D (H1) → GroC (C1a) |
| $\alpha\text{-D-Glcp-(1}\rightarrow$ | F | 4.96 → 3.95<br>F (H1) → I (H6) | 4.96 → 68.36<br>F (H1) → I (C6) |
| $\alpha\text{-D-Glcp-(1}\rightarrow$ | G | 4.96 → 3.94<br>G (H1) → M (H6) | 4.96 → 67.60<br>G (H1) → M (C6) |
| $\alpha\text{-D-Glcp-(1}\rightarrow$ | H | 4.93 → 3.82<br>K (H1) → GroC (H1a) | 4.9389 → 69.31<br>K (H1) → GroC (C1a) |
| $\rightarrow 3)\beta\text{-GalNAcp-(1}\rightarrow$ | I | 4.71 → 4.28<br>I (H1) → O (H4) | 4.71 → 75.75<br>I (H1) → O (C4) |
| $\rightarrow 3,6)\beta\text{-GalNAcp-(1}\rightarrow$ | I' | 4.69 → 4.28<br>I' (H1) → O' (H4) | 4.69 → 75.25<br>I (H1) → O (C4) |
| $\rightarrow 3,6)\beta\text{-D-GalNAcp-(1}\rightarrow$ | M | 4.58 → 4.331<br>O (H1) → A (H4) | ND |
| $\beta\text{-GlcNAcp-(1}\rightarrow$ | N | 4.54 → 3.85<br>N (H1) → J (H4) | ND |
| | | 4.54 → 3.87<br>N (H1) → C (H4) | 4.54 → 76.63<br>N (H1) → C (C4) |
| $\rightarrow 3,6)\beta\text{-GalNAcp-(1}\rightarrow$ | O | 4.52 → 3.64<br>T (H1) → GroB (H1b) | 4.52 → 72.17<br>T (H1) → GroB (C1b) |
| $\rightarrow 3)\beta\text{-GalNAcp-(1}\rightarrow$ | O' | 4.51 → 3.64<br>T (H1) → GroB (H1b) | 4.52 → 72.17<br>T (H1) → GroB (C1b) |
| $\rightarrow 3,6)\beta\text{-D-Glcp-(1}\rightarrow$ | P | 4.52 → 4.18<br>P (H1) → M (H3) | ND |
| | | ND | 4.52 → 81.84<br>P (H1) → I (C3) |
| $\rightarrow 6)\beta\text{-D-Glcp-(1}\rightarrow$ | P' | 4.50 → 4.18<br>P (H1) → M (H3) | ND |
| | | ND | 4.50 → 81.90<br>P (H1) → I' (C3) |
| $3\rightarrow)\beta\text{-D-Galp-(1}\rightarrow$ | Q | 4.48 → 4.19<br>Q (H1) → P (H6) | 4.48 → 69.31<br>Q (H1) → P (C6) |

**Supplemental Table 4:** Bacterial strains and plasmids.

| Strains/plasmids | Relevant properties/sequence | Source |
| --- | --- | --- |
| **Strains** |  |  |
| *Enterococcus faecalis* |  |  |
| OG1RF | Clinical isolate from human oral cavity | (1) |
| OG1RF Δ*epaB* | OG1RF derivative with an in-frame deletion in *epaB* | (2) |
| OG1RF Δ*epaB* + *epaB* | Complemented Δ*epaB* mutant | (2) |
| *Escherichia coli* |  |  |
| BW25113 Δ*wbbL* | wbbL deletion mutant (Keio collection) | (3) |
| **Plasmids** |  |  |
| pMV158_GFP | Plasmid constitutively expressing the Green Fluorescent Protein | (4) |
| pCV1 | Plasmid for arabinose-inducible expression of proteins in *E. coli* | (5) |
| pEpaB | pCV1-derivative for *E. faecalis* EpaB inducible expression | This work |
| pWbbL | pCV1-derivative for *E. coli* WbbL inducible expression | This work |
| pGacB | pCV1-derivative for *S. pyogenes* GacB inducible expression | (6) |

